# Distinct nanoscale architectures of GABAergic inhibitory synapses predict diverse synaptic output

**DOI:** 10.64898/2026.06.26.734878

**Authors:** Amber R. Stewart, Sara E. Gookin, Joshua D. Garcia, Kevin C. Crosby, Katharine R. Smith

**Affiliations:** Department of Pharmacology, University of Colorado School of Medicine, Anschutz Medical Campus, Aurora, CO 80045

**Keywords:** Inhibitory synapse, GABA_A_ receptor, gephyrin, nano-architecture

## Abstract

GABAergic synaptic inhibition is heterogenous across neuronal compartments, and plays a critical role in shaping local, cellular and circuit excitability. In pyramidal neurons, inhibition is mediated by GABA_A_ receptors (GABA_A_Rs) clustered at the inhibitory postsynaptic domain (iPSD). Synaptic strength depends not only on the number of GABA_A_Rs within the iPSD, but also on their precise nanoscale organization into discrete sub-synaptic domains (SSDs). These SSDs often align with presynaptic GABA release sites to form nanocolumn structures that enhance synaptic efficacy. While nanocolumn organization is increasingly recognized as a key determinant of synaptic function, most studies of GABAergic synapses have focused on archetypal dendritic synapses, which control the plasticity and integration of excitatory inputs. Nonetheless, it remains unclear whether somatic synapses - which deliver and provide robust inhibition to suppress neuronal output - share a similar nanoscale organization. Here, we used complementary super-resolution imaging approaches to directly compare inhibitory synapses in somatic and dendritic compartments. We found that somatic synapses are larger and exhibit greater structural diversity and nanoscale complexity than dendritic synapses. Dendritic synapses display relatively compact architectures with GABA_A_R SSDs frequently arranged into nanocolumns. In contrast, somatic synapses show a broader range of organizations, including aligned nanocolumns as well as more disorganized configurations with additional misaligned release sites or receptor SSDs. Computational modeling revealed that these structural differences produce distinct functional outcomes, including increased IPSC amplitude and altered kinetics at somatic synapses. Together, our findings demonstrate that nanoscale organization differentially shapes inhibitory strength and signaling properties across neuronal compartments.

**SIGNIFICANCE STATEMENT:** Diverse GABAergic synaptic inhibition is crucial to control brain excitability and its efficacy is influenced by the nanoscale trans-synaptic alignment of GABA_A_Rs and GABA release sites. Although GABA_A_R nano-architecture is defined at dendritic synapses, the extent to which this organization is conserved across GABAergic synapses with distinct synaptic properties is unknown. Using super-resolution imaging methods, we report that inhibitory synapses in the soma are larger and more structurally diverse than dendritic synapses, exhibiting both aligned and more disorganized configurations. Combined with computational modeling indicating distinct nanoarchitectures can create heterogeneous inhibitory currents, these findings suggest a key role for nanoscale organization in the generation of diverse synaptic outputs across the neuron, which could serve distinct circuit functions.

## INTRODUCTION

Inhibition in the brain is critical for regulating neural circuit function and synaptic plasticity, essential mechanisms underpinning learning, memory, and cognition (1–3). Consequently, disrupted inhibition is implicated in the pathology of numerous neurological, psychiatric, and neurodevelopmental disorders (4, 5). In the adult central nervous system, GABAergic transmission is mediated by synaptic GABA_A_ receptors (GABA_A_Rs) localized to the inhibitory postsynaptic domain (iPSD), directly opposing axonal terminals that release GABA. The number of GABA_A_Rs anchored in the iPSD directly impacts synaptic strength (6, 7), and modulating their accumulation via changes to receptor trafficking or stabilization at iPSD sites underlies key aspects of GABAergic synaptic plasticity (8). Recent studies revealed that synaptic neurotransmitter receptors, like GABA_A_Rs, are spatially organized with finer precision than previously appreciated, often forming nanoscale subsynaptic domains (SSDs) within the PSD (9, 10). For example, at forebrain excitatory synapses, AMPA receptors (AMPARs) form SSDs that align with sites of glutamate release in the presynaptic active zone, creating structures known as “nanocolumns” (11, 12). This precise AMPAR-to-release site alignment is thought to enhance synaptic efficacy since receptors are uniquely positioned for maximal binding of neurotransmitter released from presynaptic sites and likely serve as a locus for tuning synaptic strength (12–15). Similarly, we and others identified nanocolumn structures at dendritic inhibitory synapses, where synaptic GABA_A_Rs form post-synaptic SSDs that are tightly aligned with pre-synaptic GABA release sites (10, 16). Critically, disrupting this coordinated nano-alignment impairs GABAergic synaptic currents (17, 18), indicating that the subsynaptic spatial organization of GABA_A_Rs is important for efficient synaptic inhibition and thus could play a central role in shaping local dendritic excitability.

While nano-alignment appears to be a conserved feature of several synapse types (10, 11, 16, 19), variation in this architecture is observed across distinct cell-types and circuits (19–21). These differences suggest that nanoscale sub-synaptic architecture is likely shaped by cell identity, synaptic output requirements and functional demands of the circuit. Previous work investigating protein nanoarchitecture at inhibitory synapses largely focused on synapses localized to dendrites, where they provide dynamic inhibition to regulate dendritic excitability and local input integration (22, 23). However, inhibitory synapses also form across the cell soma, where they generate large-amplitude inhibitory post-synaptic currents (IPSCs) that suppress action potential firing and are critical for synchronizing circuit activity (22, 23). In agreement with these distinct functions of dendritic and somatic inhibition, ultrastructural data suggest that somatic inhibitory synapses generally possess larger pre-synaptic domains relative to those contacting dendrites (22, 24). Yet, a complete and comparative analysis of inhibitory synapse nano-architecture across the somatic and dendritic subcellular compartments has not been performed and could reveal key roles for sub-synaptic nano-organization in contributing to diverse forms of synaptic transmission within a single neuron.

In this study, we used super-resolution imaging and computational modeling to directly compare inhibitory synaptic nanoscale architecture and its predicted impact on synaptic output at synapses localized to either somatic or dendritic neuronal compartments. Our results reveal that inhibitory synapses have distinct and more heterogeneous nanoarchitectures in the soma compared with dendrites. These structural differences likely contribute to a wide range of IPSC characteristics that vary across the somatic-dendritic axis.

## RESULTS

### Inhibitory synapses localized to the soma are larger and harbor more post-synaptic SSDs than those formed on dendrites

To compare inhibitory synapse structure in somatic and dendritic sub-cellular regions, we first used three-dimensional structured illumination microscopy (3D-SIM), which enables high-throughput super-resolution imaging of synapse populations to capture their fine-scale structural features. We imaged cultured hippocampal neurons labeled with antibodies to GABA_A_Rs (surface GABA_A_R-γ2), the inhibitory post-synaptic scaffold gephyrin, and the pre-synaptic inhibitory marker VGAT (Fig. 1A), and analyzed VGAT-positive synapses from somatic and dendritic sub-cellular compartments (examples shown in Fig. 1B). Using our previously published object-based segmentation analysis (10, 17, 25, 26), we reconstructed synapses in 3D and generated volumetric data for each sub-synaptic compartment (total, pre- or post-synapse, and GABA_A_Rs or gephyrin alone: Fig. 1C). Quantification of total synapse volumes showed that inhibitory synapses formed on the soma generally had larger volumes than synapses formed on dendrites. This was the case across all synapses when analyzed together (Fig. 1D, left) as well as when comparing average synapse size within individual neurons (Fig. 1D, right). This effect was independent of whether the dendritic synapses were proximal (<50 μm) or distal (>50 μm) to the soma (SFig. 1A,B). Strikingly, somatic synapse volumes were far more heterogeneous compared to those on the dendrites: volumetric data for the somatic synapse population exhibited a wider inter-quartile range (Fig. 1D, left) and broader range of synaptic volumes compared to dendritic synapse populations (Fig.1D,E). In addition, pre- and post-synaptic compartment volumes were both larger for somatic synapses compared to dendritic synapses (Fig. 1D), indicating that both synaptic domains likely contribute to the larger and more diverse synapse volumes observed across the soma. At the iPSD, GABA_A_R compartment volumes were also significantly larger in somas compared to dendrites, suggesting more receptors are likely clustered at synapses in the soma (Fig. 1F). This difference was smaller for gephyrin and only statistically significant when paired comparisons were made between the soma and dendrites of individual neurons (Fig. 1F). Together, these results suggest the possibility that some somatic synapses with larger GABA_A_R compartment volumes may not always harbor a larger associated gephyrin scaffolding network.

**Figure 1.**
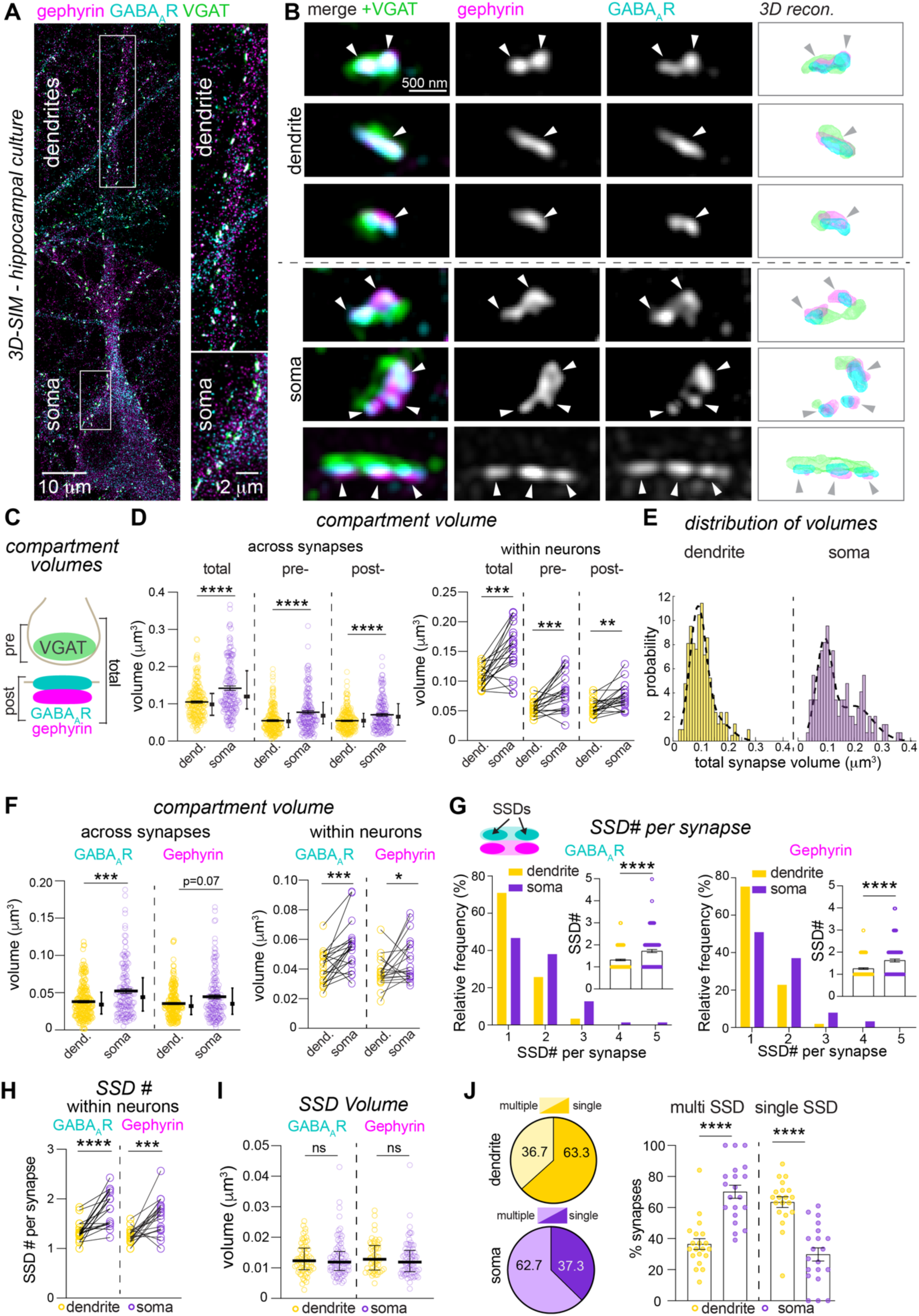
Inhibitory synapses localized to the soma are larger and harbor more post-synaptic SSDs than those formed on dendrites. (A) 3D-SIM maximum projection of hippocampal neuron in culture, labeled with antibodies to surface GABA_A_R-γ2, gephyrin and VGAT with boxes indicating example areas of the soma and dendrites. (B) Magnifications of example dendritic and somatic synapses. 3D-reconstructions of segmented images are shown on the right. (C) Summary of segmented compartment volumes generated from SIM images. (D) Quantification of total, pre- and post-synaptic volumes for dendritic and somatic inhibitory synapses across all synapses (left). Mean ± SEM is shown and median ± IQ range is shown to the right of the data points. Quantification of compartment volumes within individual neurons is shown on the right. (E) Distribution of total synapse volumes for dendritic and somatic synapses. (F) GABA_A_R and gephyrin compartment volumes for dendritic and somatic inhibitory synapses across neuronal populations (left) and within the same neuron (right). (G-H) GABA_A_R and gephyrin SSD number is increased in somatic synapses compared with dendritic synapses, across synapses (G) and within individual neurons (H). (I) GABA_A_R and gephyrin SSD volume for dendritic and somatic synapses. (J) Percentage of synapses with multiple GABA_A_R or gephyrin SSDs is increased in somatic synapses (n=7 cultures), shown across synapses (left) and averaged across neurons (right). All data: n= 210 synapses (dendrite), 150 synapses (soma), from 20 neurons, 7 independent cultures. *p ≤ 0.05, ** p ≤0.01, ***p ≤ 0.001, ****p ≤ 0.0001. Significance was assessed with Mann–Whitney (across synapses) or paired *t* test (within neurons).

We next compared the iPSD SSD characteristics between dendritic and somatic neuronal compartments. Previously, we showed that the number, but not the size of GABA_A_R and gephyrin SSDs scales-up with increasing synapse volume in dendritic iPSDs (10). As we observed that somatic synapses were much larger than dendritic synapses, we predicted that they would also contain more post-synaptic SSDs. We first quantified the mean SSD number and volume for each synapse across the dendrites and soma (Fig. 1G-I). Somatic synapses contained more numerous GABA_A_R and gephyrin SSDs compared to dendritic synapses (Fig. 1G, H) with no increase in SSD volume (Fig.1I). This finding was also supported by an increased percentage of synapses with multiple iPSD SSDs in the somatic synapse population compared to the dendritic population (Fig. 1J). We also tested whether a similar SSD scaling relationship exists in somatic synapses by drawing correlations between synapse volume and the number of associated GABA_A_R or gephyrin SSDs. Indeed, linear regression analysis showed that in both dendritic and somatic synapses, the number of SSDs increases with synapse size, and this is even more accentuated in somatic synapses (Fig. S2A,B). To confirm these results using a second independent super-resolution imaging method, we also performed direct stochastic optical reconstruction microscopy (dSTORM) of surface GABA_A_Rs (GABA_A_R- γ2) and gephyrin at VGAT-positive inhibitory synapses (Fig. S3A,B). In agreement with our SIM data, analysis of single molecule localizations and delineation of SSD structures revealed more GABA_A_R and gephyrin SSDs in somatic synapses (Fig. S3C,D). Furthermore, the mean SSD area was larger for GABA_A_Rs in somatic synapses (Fig. S3E), indicating that more GABA_A_Rs could be clustered in SSDs. Together, these results demonstrate that inhibitory synapses formed at the soma are larger, contain more SSDs of gephyrin and GABA_A_Rs, and display substantial volumetric diversity compared with their dendritic counterparts.

### Differences in dendritic and somatic synapse GABA_A_R nanoarchitecture are recapitulated in CA1 neurons in hippocampal brain slices

As synapse organization might be different in intact neuronal circuitry, we next tested the differences between dendritic and somatic synaptic nanoarchitecture in acute hippocampal slices. We optimized the imaging of GABA_A_Rs and VGAT in acute hippocampal slices by three-dimensional stimulated emission depletion (3D-STED) microscopy (Fig. 2A,B), which provides increased resolution compared to 3D-SIM and is well suited to visualizing small structures in tissue samples. We imaged inhibitory synapses in the stratum radiatum (SR) and stratum pyramidale (SP) layers of hippocampal CA1. Here, the neuropil of SR contains mostly synapses on pyramidal cell apical dendrites, and the pyramidal neuron cell bodies of the SP harbor somatic synapses (Fig. 2A,B). Analysis of volumetric synapse parameters from 3D STED images of these two regions showed that total synapse volume was larger for somatic SP synapses compared with dendritic SR synapses (Fig. 2C), with larger post-synaptic GABA_A_R and presynaptic VGAT compartments at somatic SP synapses (Fig. 2D,E). We then analyzed the SSD numbers within each synapse and found that SP synapses contained more numerous GABA_A_R SSDs than synapses in SR, suggesting a more complex nano-organization for the iPSD in somatic synapses *in situ* (Fig. 2F). Together, these analyses of inhibitory synapses in acute hippocampal slices show that the nanoscale differences we observe between dendritic and somatic synapses in hippocampal culture are mirrored *in situ*.

**Figure 2.**
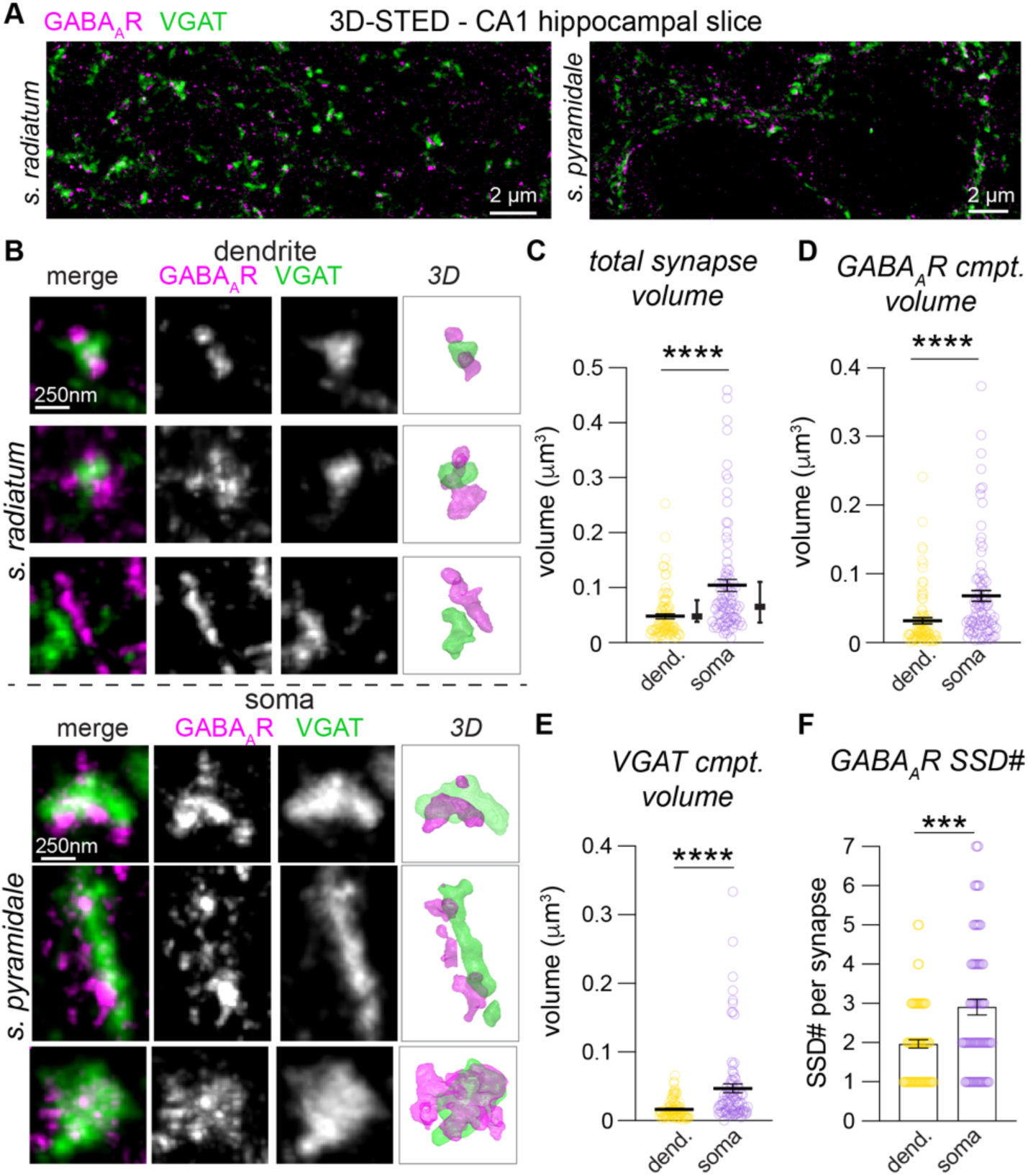
Differences in GABA_A_R nanoarchitecture in dendritic and somatic synapses are recapitulated in CA1 hippocampal neurons. (A) Maximum projected 3D-STED images of regions of area CA1 in a hippocampal slice labeled with antibodies to GABA_A_Rs and VGAT, in stratum radiatum (left) and stratum pyramidale (right). (B) 3D-STED maximum projections of dendritic and somatic synapses, labeled for GABA_A_Rs and VGAT. 3D-reconstructions of segmented images are shown on the right. (C-E) Quantification of total synapse volume (C), and GABA_A_R (D) and VGAT (E) compartment volumes. (F) Quantification of GABA_A_R SSD number per synapse in dendritic and somatic synapses. All data: n= 90 synapses (SR- dendrite), 87 synapses (SP- soma), 2 mice. ***p ≤ 0.001, ****p ≤ 0.0001. Significance was assessed with Mann–Whitney tests.

### GABA_A_R and gephyrin SSDs are less tightly associated in somatic synapses compared with dendritic synapses

As we have previously shown a robust spatial relationship between GABA_A_R and gephyrin SSDs in the iPSD of dendritic synapses (10), we next evaluated whether GABA_A_R and gephyrin SSDs are as closely associated in the soma as in the dendrites. We first generated line-scans through the iPSDs of dendritic and somatic synapses and examined synapse 3D reconstructions generated from 3D-SIM imaging (Fig. 3A). Although the majority of dendritic synapses exhibited extensive overlap between GABA_A_R and gephyrin SSDs, a subset of somatic synapses had GABA_A_R SSDs that lacked corresponding gephyrin SSDs or even a diffuse gephyrin scaffold (Fig. 3A). To quantify this observation, we calculated the 3D overlap and center-to-center distances between GABA_A_R and gephyrin SSDs in somatic iPSDs compared to those in dendrite (Fig. 3B,C). These analyses revealed reduced 3D-overlap and increased center-to-center distances between neighboring GABA_A_R and gephyrin SSDs, but only when measured from the GABA_A_R to the gephyrin SSD, suggesting that some GABA_A_R SSDs have less gephyrin in the soma (Fig. 3B,C). This finding is in line with the marginal increase in gephyrin compartment volume we observed in somatic synapses compared to dendritic synapses (Fig. 1F), and outlying populations of somatic synapses with larger GABA_A_R compartment volumes compared to their accompanying gephyrin compartments, as shown in the correlation plot in Fig. 3D. We also verified these findings through an independent method by further analysis of our dSTORM data (Fig. 3E). We quantified the area of overlap between SSDs based on the fractional area of each GABA_A_R SSD overlapping with a gephyrin SSD (as previously in (26)). This analysis revealed a similar result as with 3D-SIM: a significantly reduced fraction of GABA_A_R SSD area is overlapping with gephyrin SSDs in somatic synapses compared with dendritic synapses (Fig. 3F). Thus, our SIM and dSTORM analyses suggest that somatic inhibitory synapses tend to have more SSDs of both GABA_A_Rs and gephyrin in somatic synapses compared with dendritic synapses, there is likely a reduced association between these SSDs in at least some somatic synapses.

**Figure 3.**
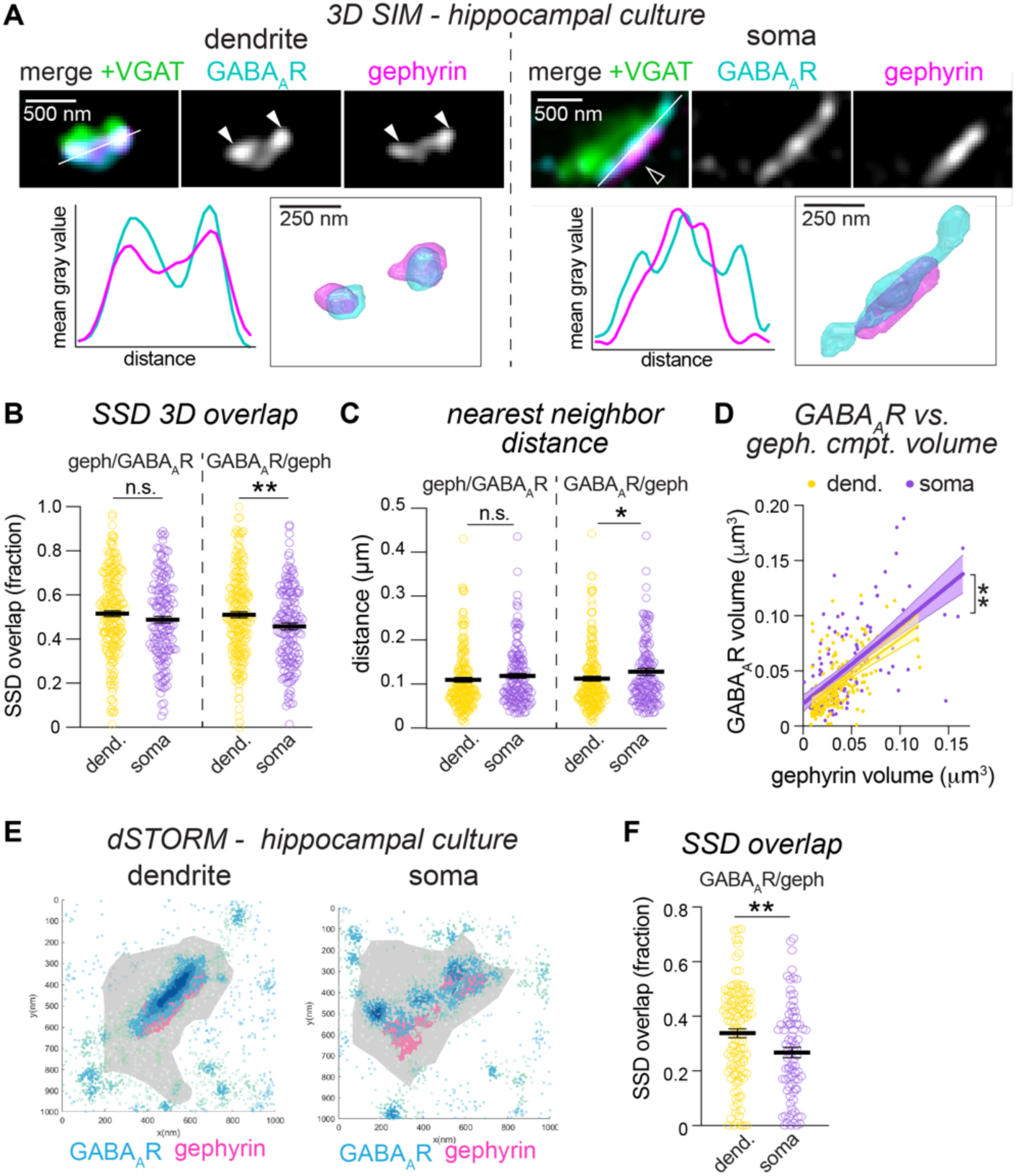
GABA_A_R and gephyrin SSDs are less tightly associated in somatic synapses compared with dendritic synapses. (A) 3D-SIM maximum projections of exemplar dendritic and somatic synapses, labeled for GABA_A_R-γ2, gephyrin, and VGAT. Closed arrowheads = overlapping SSDs, open arrowheads = non-overlapping SSDs. Line-scans are shown on the left and 3D-reconstructions of segmented images are shown on the right. (B) Quantification of 3D overlap between GABA_A_R and gephyrin SSDs. (C) Quantification of 3D nearest neighbor distance between GABA_A_R and gephyrin SSDs. (D) Correlation plot between GABA_A_R compartment volume and gephyrin compartment volume. (dend: R^2^: 0.31, soma: R^2^: 0.41, slope p = 0.68, intercept p= 0.0021). (E) dSTORM plots showing gephyrin scaffold boundary (gray) and boundaries for gephyrin SSDs (blue) with GABA_A_R SSD localizations overlaid (magenta). (F) Quantification of area overlap from dSTORM localization data. Data (A-D): n= 210 synapses (dendrite), 150 synapses (soma), from 20 neurons, 7 independent cultures. Data (E,F): n= 83 synapses (dendrite), 118 synapses (soma), from 12 neurons across 4 independent cultures. *p ≤ 0.05, ** p ≤ 0.01. Significance was assessed with Mann–Whitney test (B, C, F) or linear regression (D).

### Trans-synaptic nanoarchitecture is highly varied in the somatic sub-cellular compartment

Nanocolumnar structures are composed of pairs of post-synaptic GABA_A_R SSDs and pre-synaptic RIM1 SSDs (labeling pre-synaptic release sites (10, 27)) and are thought to contribute to synapse strength (10, 11, 17). Our previous work demonstrated that in dendrites, the number of trans-synaptic GABA_A_R-RIM1 nanocolumns scales-up with increased synapse volume (10, 17). Since somatic synapses have larger volumes and contain higher numbers of GABA_A_R SSDs, we hypothesized that these synapses may also contain more numerous nanocolumn structures than dendritic synapses, which could contribute to stronger synapses. In these experiments we used STED microscopy to imaged surface GABA_A_Rs and predicted sites of pre-synaptic GABA release sites by imaging RIM1 (10, 11), while broadly delineating the iPSD by labeling gephyrin. Fig. 4A shows an overview STED image of a representative hippocampal neuron, that highlights the dendritic and somatic regions (on right). Representative STED images and 3D-reconstructions of individual dendritic and somatic synapses are depicted in Fig. 4B. We performed a similar segmentation analysis as for 3D-SIM with minor modifications (as in (17)), and quantified compartment volumes, SSD volumes and SSD numbers for GABA_A_Rs and RIM1 for each synapse. Notably, the RIM1 compartment was larger in somatic synapses (Fig. 4C), and contained higher numbers of RIM1 SSDs (Fig. 4D), which is suggestive of a more expansive active zone (AZ) and potentially more numerous GABA release sites in this synapse population. In agreement with our SIM data (Fig. 1), GABA_A_R compartment volume in somatic synapses imaged using STED was also larger and contained more SSDs than dendritic synapses (Fig. 4E,F). Interestingly, the use of higher-resolution STED microscopy also revealed that individual GABA_A_R and RIM SSD volumes were larger in somatic synapses (Fig. S4A). This effect was particularly striking for GABA_A_R SSDs, and similar to our dSTORM results (SFig. 2), suggesting more receptors are organized into individual SSDs in somatic synapses than in dendritic synapses.

**Figure 4.**
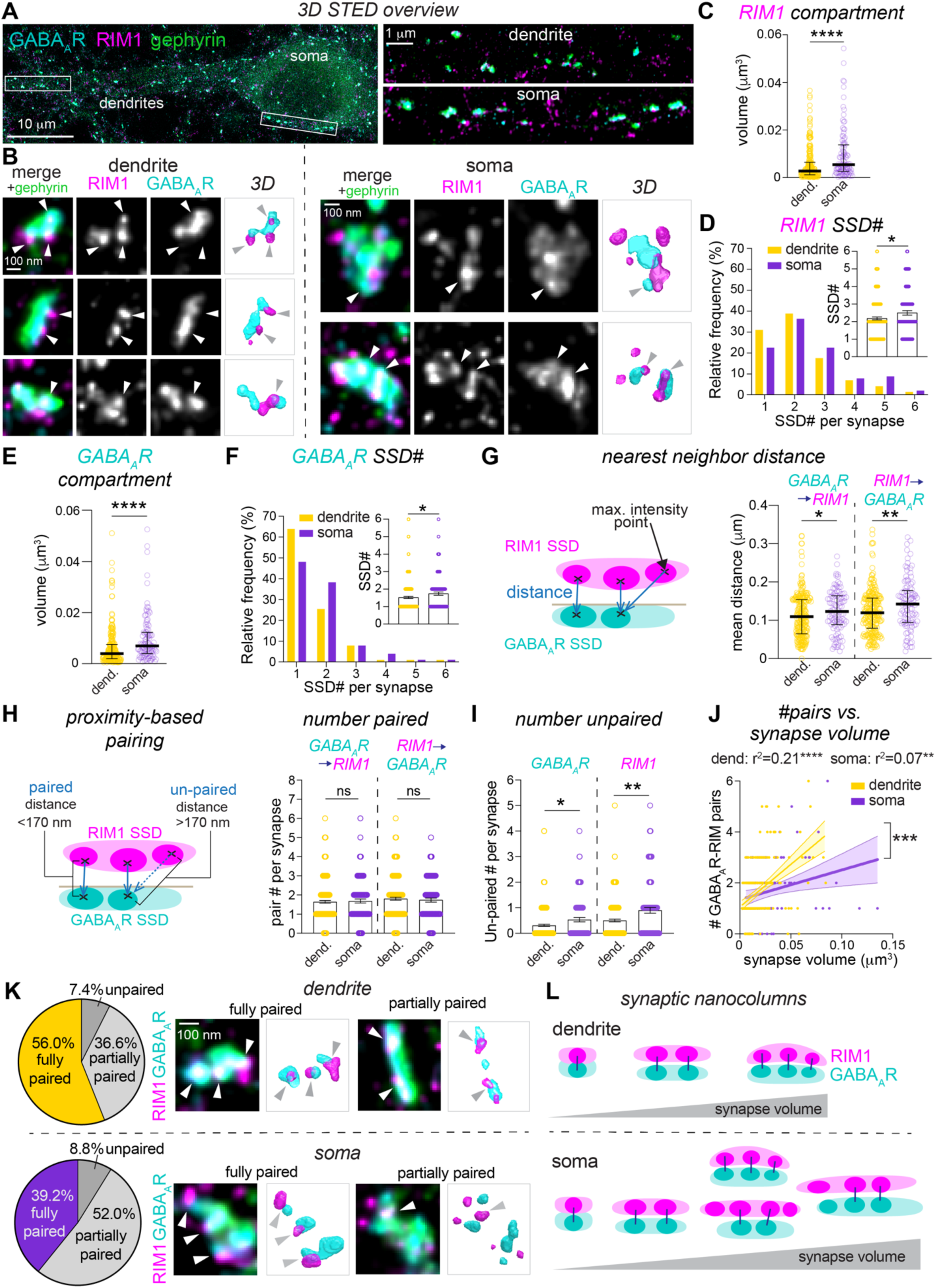
Trans-synaptic nano-architecture is highly varied in the somatic sub-cellular compartment. (A) Maximum projected 3D-STED image of cultured hippocampal neuron labeled with antibodies to RIM1, GABA_A_R-γ2, and VGAT (left). Boxes indicate example regions of soma and dendrites shown on the right. (B) Magnifications of example dendritic and somatic synapses from 3D-STED images with 3D-reconstructions. Arrowheads= GABA_A_R and RIM1 pairs. (C) RIM1 compartment volume determined by 3D-STED. (D) Frequency distribution and quantification of RIM1 SSD number per synapse. (E) GABA_A_R compartment volume determined by 3D-STED. (F) Frequency distribution and quantification of GABA_A_R SSD number per synapse. (G) Quantification of inter-SSD nearest neighbor 3D distance per synapse. (H) Quantification of number of pairs per synapse, determined by the nearest neighbor distance (I) Quantification of number of unpaired SSDs per synapse, determined by the nearest neighbor distance. (J) Linear correlation between number of GABA_A_R-RIM1 nanocolumns and total synapse volume for dendritic and somatic synapses; (dend: R^2^= 0.21, =p<0.0001; soma: R^2^= 0.069, p= 0.074; slope p= 0.0005). (K) Quantification of the percentage of dendritic and somatic synapses that have fully, partial or no pairing between GABA_A_R and RIM1 SSDs. Additional synapse examples are shown on the right (merged and 3D reconstructed images are shown). (L) Cartoon summarizing the trans-synaptic architecture of dendritic and somatic synapses as it changes with synapse volume. Data in C-K: n= 216 synapses (dendrite), n= 102 synapses (soma), from 11 neurons, 3 independent cultures. *p ≤ 0.05, ** p ≤ 0.01, ***p ≤ 0.001, ****p ≤ 0.0001. Significance was assessed with Mann–Whitney test (C-I) and linear regression (J).

To determine whether the more numerous GABA_A_R and RIM1 SSDs in somatic synapses formed more trans-synaptic nanocolumns, we adapted our proximity-based distance analysis (10), to analyze STED images. We first calculated the nearest neighbor distance in 3D between maximum intensity points of the fluorescent signal in each RIM1 SSD and its nearest neighboring GABA_A_R SSD (and vice versa: Fig. 4G (17)). Surprisingly, this distance was significantly increased in the somatic synapse population, irrespective of whether we measured from RIM to GABA_A_Rs or GABA_A_Rs to RIM (Fig. 4G). This result indicates increased separation between GABA_A_R and RIM SSDs in somatic synapses compared with dendritic synapses and could reflect differences in pre- and post-synaptic nanoscale alignment. To test the extent of association between GABA_A_R and RIM1 SSDs in forming a nanocolumn, we classified whether GABA_A_R SSDs were ‘paired’ to their nearest RIM1 SSD, based on their proximity to each other (Fig. 4H). We then used a pre-determined 3D distance to classify SSDs as paired or un-paired with a nearest neighboring SSD (determined by heuristic assessment of 3D reconstructions and comparisons between experimental data and simulated control data (10, 17): Fig. 3H, see methods for details). Inter-SSD distances greater than 170 nm were categorized as ‘unpaired’, whereas those that were shorter than 170 nm were categorized as ‘paired’ and thus defined as a nanocolumn. Surprisingly, even though somatic synapses contained more SSDs, the number of trans-synaptic pairs was similar in dendritic and somatic synapses (Fig. 4H). Indeed, quantification of the unpaired GABA_A_R or RIM SSDs in each synapse revealed a significant increase in un-paired or ‘orphaned’ SSDs in somatic synapses (Fig. 4I, SFig. 4B). In agreement with this result, we found a reduced percentage of synapses with full pairing between GABA_A_R and RIM1 SSDs in the soma compared with the dendrites, and an increased percentage of partially paired synapses (Fig. 4K: additional example synapses are shown). The observation of unpaired SSDs was significant for both GABA_A_R and RIM1 SSDs, but the difference was much larger for RIM1, suggesting that RIM1 SSDs are more often unpaired compared to GABA_A_R SSDs. As we found previously (10), in dendritic synapses the number of nanocolumns per synapse scaled-up with synapse size (Fig. 4J). However, this correlation was markedly reduced in the somatic synapse population (Fig. 4J), indicating that larger somatic synapses do not necessarily harbor higher numbers of nanocolumns and may follow distinct nano-architectural rules. Together, these data show that somatic synapses have a distinct and highly varied transsynaptic nanoarchitecture compared with dendritic synapses; this organization is characterized by more numerous RIM SSDs, some of which lack an associated post-synaptic GABA_A_R SSD, and a more disorganized and heterogeneous topography.

### Simulations reveal that different trans-synaptic nanoarchitectures may contribute to diverse synapse outputs

Acute disruption of GABA_A_R nano-alignment with pre-synaptic release sites reduces synaptic strength (17, 18), indicating that trans-synaptic nanocolumn structures contribute to the efficacy of inhibitory synapse function. Yet here we observed striking nano-structural heterogeneity of inhibitory synaptic architecture across the somato-dendritic axes of hippocampal neurons, suggesting that different nanoscale architectures could potentially impact the output of an inhibitory synapse. To test whether the different nano-organizations we observed across the somatic and dendritic compartments could contribute to synaptic output, we employed a computational approach to simulate post-synaptic inhibitory currents (IPSCs) that could be produced by the distinct nanocolumnar arrangements we observed in our experiments. We performed Monte-Carlo based stochastic simulations of GABA release, its binding to GABA_A_Rs, receptor opening and the generation of an IPSC. We adopted previously published synapse geometries and reaction-diffusion elements for the inhibitory synapse (28, 29), and a gating and kinetic scheme for synaptic GABA_A_Rs (30) with minor modifications (see methods). Our STED dataset provided the positioning of pre- and post-synaptic elements relative to each other, relative SSD sizes and numbers per synapse (Fig. 5A). To validate our model, we first ran simulations for an archetypal dendritic synapse with two release sites, modeling multi-vesicular release from both sites. A characteristic IPSC was generated (Fig. 5B,C), with a mean peak amplitude of 13.3 pA and fast activation and decay kinetics, which were reflected in a short time to peak amplitude and time to decay half. However, either positioning the post-synaptic GABA_A_R SSDs away from the release sites (Fig. 5D) or arranging the same number of receptors uniformly throughout the iPSD substantially reduced the IPSC amplitude and produced significantly slower current kinetics than when GABA_A_R were clustered directly opposite release sites (Fig. 5E). Critically, these results support previous electrophysiological experiments showing that acute disruption of GABA_A_R-RIM association reduces IPSC strength (17, 18) and indicate that our model generates IPSCs that are disrupted by perturbations in the inhibitory nanocolumn architecture.

**Figure 5.**
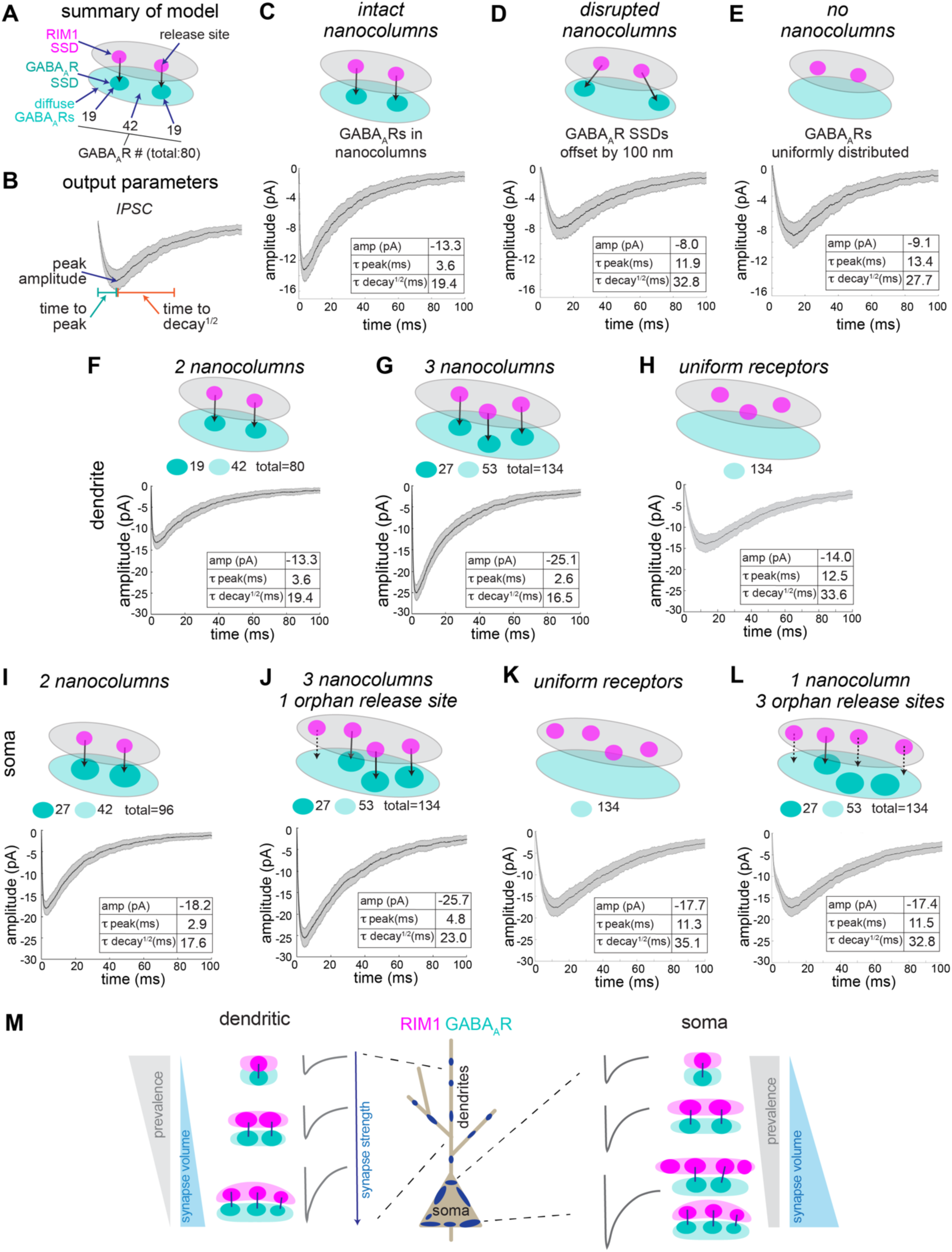
Computational simulations reveal that different inhibitory synaptic nanoarchitectures may contribute to diverse synaptic outputs. (A) Schematic showing positions of GABA_A_R SSDs and presynaptic GABA release sites used in simulation. 3D spatial architecture was based on STED data, the geometric components of the three model synapses were the same except for the presence and/or alignment of the GABA_A_R SSDs. Receptor numbers are those used in simulations C-E. For all simulations, 500 GABA molecules were released from each release site. Traces are 160 runs; the black trace represents the mean value, and gray represents the standard deviation. Tables show peak amplitude, time to peak and time to decay^1/2^. (B) IPSC parameter measurements generate from simulation; peak amplitude, time to peak and time to decay^1/2^, shown in table for each simulation. (C-E) Simulations of IPSCs from synapses with: (C) two nanocolumns, (D) disrupted nanocolumns, or (E) where receptors are uniformly distributed. (F-H) Simulations of IPSCs from dendritic synapses with: (F) 2 nanocolumns (same as C), (G) 3 nanocolumns in a larger synapse with more receptors, and (H) diffuse receptors in a larger synapse. (I-L) Simulations of IPSCs from somatic synapses with: (I) 2 nanocolumns with larger post-synaptic SSDs (containing more receptors), (J) more release sites in a larger synapse with 3 nanocolumns and an orphaned release site, (K) more release sites in a larger synapse with 1 nanocolumn and 3 orphaned release sites, (L) diffuse receptors in a larger synapse. (M) Summary cartoon of inhibitory nanocolumn structure and predicted IPSCs across somatic and dendritic regions.

Next, we used this model to evaluate the properties of IPSCs generated by the different synaptic nanoarchitectures observed in the dendritic compartment. In the dendrites there is a strong correlation between synapse volume and the number of nanocolumns within the synapse (Fig. 5J;(10)), which suggests that more nanocolumns could contribute to synapse strength. Thus, we compared the IPSCs produced by synapses with either 2 nanocolumns (Fig. 5F), or with more receptors in a larger synapse organized into 3 nanocolumns (Fig. 5G). The addition of a third nanocolumn incorporating more receptors significantly increased the amplitude of the simulated IPSC, with similar kinetic properties to the condition with 2 nanocolumns (Fig. 5F). Crucially, this effect was only apparent with the addition of GABA_A_Rs clustered into an SSD positioned opposite a putative release site, as we observed very little increase in amplitude if the same number of receptors were added uniformly to the iPSD (Fig. 5H). This result demonstrates that in dendrites, the observed scaling-up of nanocolumn number with synapse size is likely correlated with increased synapse strength.

We then simulated the IPSCs that may be generated by nanoscale organizations detected in the somatic compartment. First, we modeled a commonly observed synaptic cartography found in the soma: two nanocolumns, but larger GABA_A_R SSDs than in the dendrites (that we assumed would contain more GABA_A_Rs: Fig. 5I). These somatic synapses were often of a similar size as dendritic synapses, but exhibited increased IPSC amplitude (even with the same nanocolumnar structure), presumably due to the positioning of more GABA_A_Rs in the nanocolumn near to sites of GABA release. Some of the larger somatic synapses had the key feature of more numerous and often ‘orphaned’ RIM SSDs, suggesting sites of GABA release without a post-synaptic GABA_A_R SSD. To assess the impact of orphaned release sites on somatic IPSCs, we examined a large somatic synapse with 3 nanocolumns, but 1 orphan release site (see Fig. 4B). This architecture produced a large amplitude IPSC, similar to larger dendritic synapses (Fig. 5G) and significantly larger than the control condition where receptors were randomly distributed in the iPSD (Fig. 5K). Yet, this IPSC exhibited substantially slower decay kinetics than found in the dendritic simulations, suggesting the presence of an unpaired release site can lengthen the time to full receptor activation and the decay profile of the IPSC. A second example showed that more orphaned release sites slowed the IPSC kinetics even further but also reduced IPSC amplitude. This result suggests that GABA release from un-paired release sites likely enables a longer period of receptor activation and slows the post-synaptic current, suggesting that this nanoarchitecture could create IPSCs with diverse temporal characteristics across somatic synapses. Thus together, these simulation data illustrate that the nanoscale architectures we observe experimentally may produce IPSCs with distinct features and generate diversity in synaptic inhibition across the somato-dendritic axis.

## DISCUSSION

In this study we have used complimentary super-resolution microscopy approaches to directly compare the nanoscale architecture of inhibitory synapses formed onto dendritic and somatic subcellular compartments. We show that synapses formed on the soma are not only consistently larger than those formed on dendrites, but are also more heterogeneous, both in terms of their volume and the complexity of their nanoscale sub-synaptic architecture. Our results indicate that distinct nanoscale principles exist for inhibitory synapses, which depend on the sub-cellular context of the synapse and our results suggest a new mechanism where synaptic structural diversity could underlie heterogeneity in synapse function. The wide variety of nano-organizations that we observed across the somato-dendritic axis suggests that the nanoscale organizational principles for proteins at synapses may be far more varied than first thought. This notion agrees with recent work indicating that nano-architectural motifs at glutamatergic synapses could be both cell-type and circuit specific (20, 21). Thus, the nano-organization of synapses is likely an important contributing factor to their diverse functions and act as a mechanism to enable different types of transmission to serve distinct circuit requirements.

Our detailed analysis of synapse volume showed that somatic synapses were consistently larger than dendritic synapses, which aligns with prior studies using light and serial electron microscopy demonstrating that inhibitory terminals contacting the soma have larger areas than those innervating dendrites (22, 24). Interestingly, this difference in pre-synaptic area still applies when presynaptic boutons from the same interneuron innervate both the dendrites and soma (24, 31), indicating that the size of the presynaptic terminal could be strongly dependent on properties and location of the post-synaptic compartment. This idea is reinforced by a strong linear relationship between GABAergic terminal area and the size of the post-synaptic target (31). For instance, thinner dendrites tend to be innervated by smaller synaptic terminals whereas thicker dendrites are usually associated with larger synaptic inputs (31). Functionally, this relationship is in line with dendrites requiring less hyperpolarization to shunt and regulate local dendritic excitability compared with the larger soma that likely requires greater inhibitory strength to effectively influence membrane excitability. Unexpectedly, somatic synapse volumes were considerably more heterogeneous compared to their dendritic counterparts. This variability may partly arise from the larger surface area of the somatic compartment, which could permit a broader range of synapse sizes, including very large synapses. In contrast, dendritic synapses may be limited in their maximum size by the spatial constraints of the dendritic shaft, resulting in a narrower range of possible synaptic volumes.

In dendrites, we have demonstrated that post-synaptic SSDs serve as ‘building blocks’ that scale in number as synapses increase in volume (10). Here, we observed a similar correlation in the dendrites, suggesting that this building block principle may also operate in somatic synapses and could even be more pronounced in this compartment. Thus, the nanoscale architecture of the iPSD appears to follow similar organizational principles in the soma and dendrites, which is reminiscent of glutamatergic synapses, where modules of glutamate receptors and scaffolds are shown to be added to synapses in dendritic spines as they grow (32). Although the GABA_A_R compartment volume was consistently larger in somatic synapses, this was not the case for the gephyrin scaffold, which on average had a similar volume between dendritic and somatic compartments. This observation suggests that the relationship between GABA_A_Rs and gephyrin may be different in somatic synapses. Indeed, SIM and STORM image analysis revealed a reduced spatial association between GABA_A_R and gephyrin SSDs, indicating that at some somatic GABA_A_R SSDs are less associated with, or even lack, the gephyrin scaffold. This finding supports the existence of alternative mechanisms that maintain the nanoscale clustering of GABA_A_Rs. Such mechanisms could involve the dystroglycan complex (DGC (33)), the scaffolding proteins GIT1 (34) and S-CAM (35), or the GABA_A_R accessory protein LHFPL4/GARLH (36, 37).

Given their larger volume and greater number of post-synaptic GABA_A_R SSDs, we initially hypothesized that somatic synapses would contain more trans-synaptic nanocolumn structures than dendritic synapses, which could contribute to increased synapse strength. Although somatic synapses contained more RIM1 and GABA_A_R SSDs, this did not consistently translate into greater numbers of nanocolumn pairs. Instead, we observed highly heterogeneous nanocolumn organizations throughout the somatic synapse population, with some synapses containing multiple nanocolumns while others contained a mixture of nanocolumns together with extraneous, ‘orphaned’ RIM SSDs without a corresponding post-synaptic GABA_A_R SSD. The functional role of these orphaned RIM SSDs observed here remains unclear, but it is possible that they could be rapidly recruited to form new nanocolumns during activity-dependent synaptic plasticity, without needing to add new material to the presynaptic terminal. Alternatively, expanded AZs could also support multi-vesicular release and lead to higher concentrations of GABA in the synaptic cleft (38), raising the prospect that some somatic synapses might operate effectively without strict nano-alignment. Indeed, distinct patterns of activity and modes of presynaptic GABA release which lead to increased concentrations of GABA in the synaptic cleft could override a requirement for precise alignment of pre- and post-synaptic nanoscale organization at some synapses. For example, somatic-innervating PV basket cells have fast-spiking firing patterns, while SST-expressing dendritic innervating interneurons generally have a reduced spiking output (39), which could result in different concentrations of GABA in the synaptic cleft. Furthermore, asynchronous release that is prevalent at synapses innervated by CCK interneurons could prolong the concentration of GABA in the synaptic cleft over longer periods of time (40). Thus, it is possible that at these different synapses GABA_A_Rs may not need to be positioned directly near release sites to be efficiently activated. Another related factor that could influence the requirement for precise nanoarchitecture is the degree of GABA_A_R occupancy at each synapse, which is highly variable both between different cell types and also within individual neurons (41, 42), and is thought to be dependent on the post-synaptic target cell (41). If receptor saturation occurs at a particular synapse, nanoscale alignment between release sites and receptors may be less important. In contrast, under non-saturating conditions, the optimal positioning of the receptor near sites of release could be critical. Thus, it is likely that in addition to the sub-cellular compartment in which the synapse resides, a combination of GABA release properties and synapse receptor occupancy contribute to the requirement for a precise nanoarchitecture for inhibitory synaptic function.

The molecular complexity of inhibitory pre- and post-synaptic domains is striking, and it is highly likely that different synaptic organizing complexes across different regions of the neuron could create distinct nanoarchitectures in inhibitory synapses. Based on previous work, post-synaptic neuroligin-2 (NL2) is likely a key player in creating nanocolumns (18, 26) through interactions with pre-synaptic neurexins (43). However, it is likely that other inhibitory synapse adhesion systems are also important for nanocolumn formation, including neuroligin-3 – Nrxn and DGC – Nrxn (44, 45). NL2 also participates in multiple different iPSD complexes which could contribute to or modulate nanocolumn generation (including MDGAs (46, 47), collybistin (48), GARLH/LHFPL4 (36, 37) and SCAM (49)). Thus, the diversity in synaptic nanoarchitecture could be generated through the variety of these NL2/NL3 post-synaptic interactors and transsynaptic interactions with distinct and alternatively spliced presynaptic Nrxns (43, 50). In addition, multiple GABA_A_R subtypes are expressed in a single CA1 pyramidal neuron, conferring a range of distinct pharmacological and functional properties to synaptic function (51, 52). Although some GABA_A_R subtypes are reliably shown to be at specific synapse types (53, 54), synapse-specificity for the most common receptor subunits is still unclear (55–57). It is therefore tempting to hypothesize that different subtypes could have different subsynaptic positioning depending on affinity for GABA and the requirements of a particular synapse. Future work will need to combine our knowledge of distinct proteins at inhibitory synapses with nano-architectural observations, along with functional readouts, to determine the precise contribution of different synaptic molecules in the formation, maintenance and modulation of nanocolumns at inhibitory synapses.

A computational modeling approach enabled us to make several important conclusions regarding the role of nanocolumns in inhibitory synaptic function. Firstly, our simulations show that moving the post-synaptic GABA_A_R SSDs away from the pre-synaptic release sites substantially reduced IPSC amplitude to similar levels as when the receptors are uniformly distributed in the iPSD. This result indicates that the precise positioning of GABA_A_Rs opposite release sites is an important contributor to synaptic efficacy and complements experimental evidence showing that acute perturbation of the inhibitory nanocolumn by moving GABA_A_Rs away from release sites results in reduced IPSC amplitudes (17, 18). Secondly, in dendritic synapses we find that merely increasing the number of GABA_A_Rs randomly into the iPSD is not sufficient to increase synaptic strength; instead, the receptors require incorporation into SSDs opposite a release site to increase IPSC amplitude. This result supports past experiments showing that recruitment of AMPARs randomly to the excitatory PSD does *not* increase EPSC amplitude (14), suggesting that newly added AMPARs need to be accurately positioned opposite glutamate release sites to increase synaptic strength. This result suggests that GABA_A_Rs may follow a similar rule, and receptors that are recruited to the synapse during plasticity, such as during inhibitory long-term potentiation (58), may need to be positioned into existing or newly made nanocolumns to increase IPSC amplitude. Our simulations of somatic synapse cartography showed that the broad range of nanoarchitectures observed on the somatic compartment generated IPSCs with a wide range of characteristics. For example, the addition of an extra orphaned release site to the presynaptic domain slowed the IPSC time-course, producing a longer time to peak amplitude and prolonged decay. The impact of this synaptic nano-organization on IPSC kinetics is particularly interesting because IPSC time courses are highly diverse even within a single neuron and are essential for encoding different forms of inhibition that underly neuronal information processing (52). It is possible that this kind of mechanism could function alongside differential GABA_A_R sub-type expression to generate diversity of IPSC kinetics across a single neuron (52).

Together, this work indicates that the nanoscale organization of GABAergic inhibitory synapses can vary greatly within a single neuron and may contribute to the diversity of synaptic inhibition across the somato-dendritic axis. Our findings stress the importance of sub-synaptic nanoscale organization of GABA_A_Rs for the efficacy of inhibitory synaptic strength and indicate that their architecture is likely a locus for generating different forms of synaptic inhibition. Going forward it will be essential to determine how different forms of synaptic inhibition are impacted by nanocolumn perturbation and to decipher how distinct interneuron sub-types and GABAergic molecular components control the patterning of nanoscale architecture at inhibitory synapses. Furthermore, understanding how nano-architectural organization of somatic and dendritic inhibitory synapses changes in various forms of inhibitory plasticity and models of disease where inhibition is impaired, will shed light on new mechanisms for modulating inhibitory synapse function.

## MATERIALS AND METHODS

Detailed descriptions of primary neuron culture, immunocytochemistry, immunohistochemistry, microscopy, image analysis, computational modeling and statistics can be found in *SI Appendix, Materials and Methods*.

### Dissociated Hippocampal Cultures

All animal procedures were approved by the Institutional Animal Care and Use Committee at the University of Colorado Anschutz Medical Campus. primary neuronal cultures were prepared from dissected hippocampi of mixed sex neonatal rat pups (postnatal day 0-1) as previously described (10), and plated on poly-D-lysine coated #1.5 glass coverslips. Cultures were maintained at 37°C, 5% CO_2_ for 15-18 days prior to experiments.

### Immunocytochemistry (ICC)

Cultures were fixed in 4% PFA + 4% sucrose in PBS for 5 min at RT, washed and blocked in 5% BSA + 2% Normal Goat Serum (NGS) in PBS for 30 min. Neurons were incubated with primary antibodies to surface GABA_A_R-γ2 under non-permeabilized conditions for 1 h at RT. Cells were washed permeabilized in 0.5% NP-40 for 2 min and blocked for 30 min. Gephyrin 3B11, VGAT, and RIM1 staining was performed with primary antibodies in block for 1 h. Coverslips were washed and incubated with appropriate secondary antibodies for 1 hr (for 3D-SIM, with Alexa dyes) or overnight (for STED, with Abberior STAR dyes). Coverslips were washed and mounted with ProLong Gold. Further details for ICC and immunohistochemistry can be found in the SI Materials and Methods.

### Image Acquisition

3D-SIM images were acquired with a Nikon SIM-E Structured Illumination super-resolution microscope with a 100x/1.49 NA objective, 488, 561, and 640 laser diodes and an ORCA-Flash 4.0 sCMOS camera (Hamamatsu). 3D-STED images in neuron culture were acquired using an Abberior STEDYCON addition to an Olympus confocal microscope equipped with a 100X/1.45 NA objective, 4 excitation lasers and 4 corresponding single-photon counting APD detectors (avalanche photodiodes), and a 775nm laser for stimulated emission depletion. For imaging in hippocampal CA1 brain slices, an Abberior Infinity 3D-STED microscope was used. dSTORM was performed on a Zeiss Elyra P.1 TIRF microscope using a Zeiss Plan Apochromat TIRF 100x/1.6 NA oil objective, a quad-band dichroic (405/488/561/642), iXon+ EMCCD camera (Andor). See *SI Appendix, Materials and Methods* for full details of microscopy methods.

### Super-resolution imaging analysis

3D-SIM and STED images were analyzed using MOSAIC segmentation analysis with FIJI and Matlab (10, 17, 25). dSTORM acquisitions were analyzed as in (25, 26). See *SI Appendix, Materials and Methods* for full details of microscopy analysis.

### Computational Modeling of Inhibitory Synaptic Currents

Monte-Carlo simulations were performed using the mCell platform (version 4.0.6) (59, 60). Model geometries based on our STED/SIM data, were constructed in the CellBlender (version 2.93) module of mCell. Full details of simulations can be found in *SI Materials and Methods*.

## ACKNOWLEDGEMENTS

This work was supported by R01MH119154 and R01MH128199 to K.R.S. A.R.S. was supported by Ruth L. Kirschstein individual predoctoral National Research Service Award (NRSA) F31NS141320 and T32GM007635. J.D.G. was supported by an NIH Blueprint Diversity Specialized Predoctoral to Postdoctoral Advancement in Neuroscience (D-SPAN) Award (FNS120640A). We thank members of the Smith Lab, and Drs. Mark Dell’Acqua, Matthew Kennedy and David DiGregorio for helpful discussions. Contents are the authors’ sole responsibility and do not necessarily represent official NIH views. STORM and STED Imaging was performed in the Advanced Light Microscopy Core (ALMC) facility of the NeuroTechnology Center at the University of Colorado Anschutz Medical Campus, which is supported in part by Diabetes Research Center Grant (P30 DK116073). We would like to thank Dominik Stich from the ALMC for assistance with the STED and dSTORM imaging.

## Notes

### Competing Interest Statement

The authors have declared no competing interest.

