## Supplemental Figures for "Distinct nanoscale architectures of GABAergic inhibitory synapses predict diverse synaptic output"

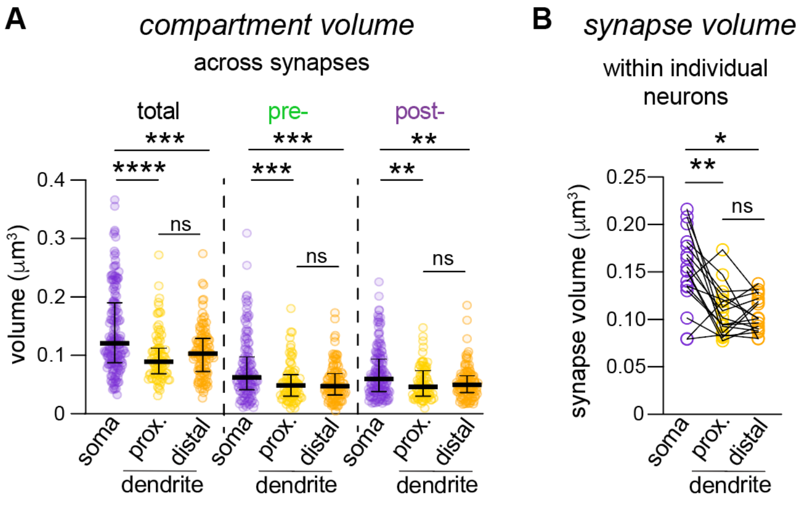


**Supplemental Figure 1. Further analysis of compartment volumes by 3D SIM.**
(A) Quantification of total, pre- and post-synaptic compartment volumes in proximal (<50µm from soma) and distal (>50µm from soma) segments of neuronal dendrites.
(B) Quantification of total synapse volume in somatic and proximal/distal dendritic synapses within individual neurons.

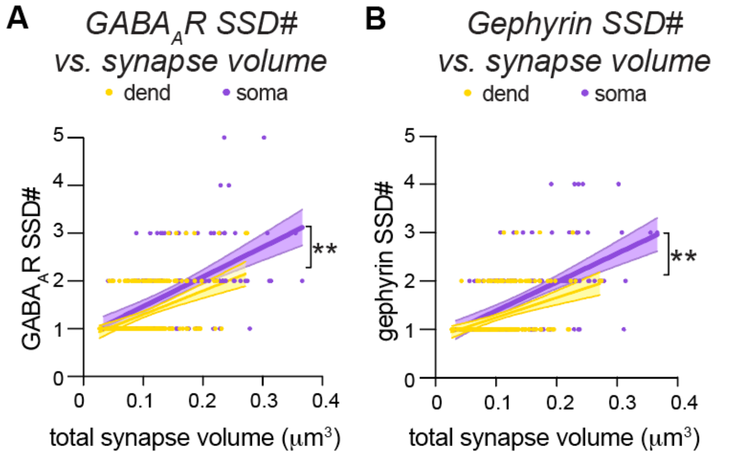


**Supplemental Figure 2. Additional correlation analysis of SIM images.**
(A) Correlation plot between $GABA_AR$ SSD number and total synapse volume. (dend: $R^2$: 0.17, soma: $R^2$: 0.29, slope p = 0.27, intercept p= 0.0045).
(B) Correlation plot between gephyrin SSD number and total synapse volume. (dend: $R^2$: 0.14, soma: $R^2$: 0.30, slope p = 0.077, intercept p= 0.0051).

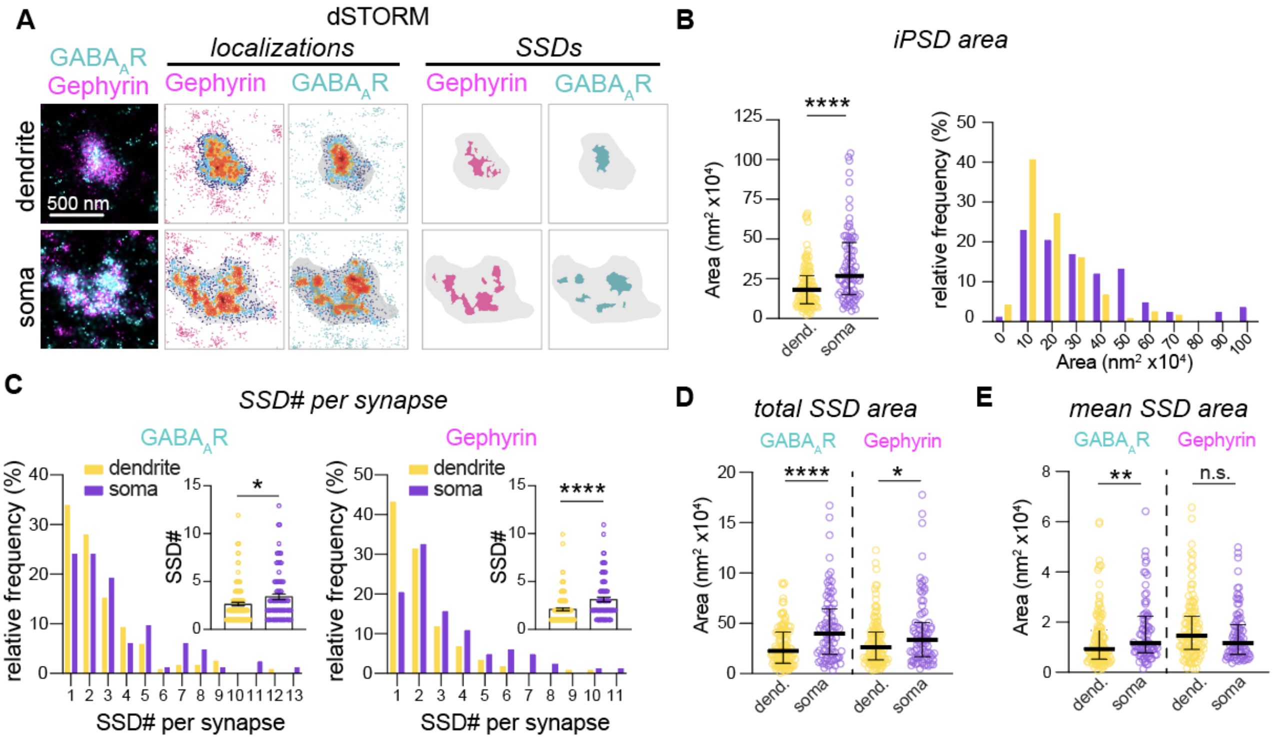


**Supplemental Figure 3. Additional analysis of GABAergic iPSDs by dSTORM.**
(A) dSTORM rendered images of inhibitory synapses labeled with antibodies to GABA$_A$R-$\gamma$2 and gephyrin. Localization plots of GABA$_A$R-$\gamma$2 and gephyrin dSTORM data with warmer colors indicating higher densities of localizations and the grey boundary representing the gephyrin post-synaptic domain. Individual SSDs are shown in binary representations of higher density regions for GABA$_A$R and gephyrin, again with the grey boundary representing the gephyrin post-synaptic domain.
(B) Quantification and frequency distribution of iPSD area, derived from gephyrin localizations. $p<0.0001$.
(C) Frequency distribution and quantification of GABA$_A$R and gephyrin SSD number for dendritic and somatic synapses determined by dSTORM. GABA$_A$Rs: $p= 0.03$, gephyrin: $p<0.0001$.
(D) Quantification of total combined SSD area. GABA$_A$Rs: $p<0.0001$, gephyrin: $p=0.04$.
(E) Quantification of mean SSD area per synapse. GABA$_A$Rs: $p=0.01$, gephyrin: $p=0.11$.

For all analyses: n= 83 synapses (dendritic), 118 synapses (somatic), from 12 neurons across 4 independent cultures.

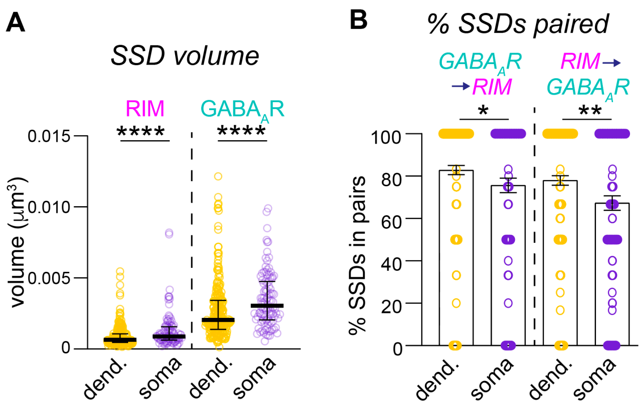


**Supplemental Figure 4.** Additional analysis of RIM and $GABA_A$Rs by STED.
(A) Quantification of SSD volume for RIM and $GABA_A$Rs.
(B) Quantification of % SSDs paired per synapse.
