## Supplemental materials and methods for "Distinct nanoscale architectures of GABAergic inhibitory synapses predict diverse synaptic output"

### SI MATERIALS AND METHODS

#### Dissociated Hippocampal Cultures

All animal procedures are in accordance with the National Institutes of Health (NIH) *Guide for the Care and Use of Laboratory Animals* and approved with the Institutional Animal Care and Use Committee at the University of Colorado, Anschutz Medical Campus. As previously described (1, 2), primary neuronal cultures were prepared from dissected hippocampi of mixed sex neonatal rat pups (postnatal day 0-1) and dissociated in papain. Neurons were seeded in MEM and 10% FBS containing penicillin/streptomycin onto 18mm #1.5 glass coverslips at a density of 150,000-200,000 cells coated with poly-D-lysine. MEM was replaced 24h following plating with Neurobasal-A (NB) medium (GIBCO) supplemented with B27 (GIBCO) and 2 mM GlutaMAX (Gibco). Feedings were performed every 5 days by removing half the media and replacing with new NB media. To limit growth of actively dividing cells, mitotic inhibitors (uridine fluoro deoxyuridine) were added at day 5. Cultures were maintained at 37°C, 5% $CO_2$ for 15-18 days prior to experiments.

#### Immunocytochemistry (ICC)

Neuronal cultures grown on coverslips were fixed in 4% PFA solution (4% sucrose, 1X PBS and 50mM HEPES (pH 7.5)) for 5 min at room temperature (RT) followed by three 5 min washes with 1X PBS. Coverslips were then blocked in 5% BSA, 2% Normal Goat Serum (NGS) and 1X PBS at RT for 30 min. Surface $GABA_AR$-$\gamma$2 (1:500 Synaptic Systems Guinea Pig – 224004) staining was performed under nonpermeabilized conditions in blocking solution for 1 hour at RT. Coverslips were washed for 5 min (3X) in PBS followed by permeabilization in 0.5% NP-40 for two min and blocked at RT for 30 min. Gephyrin 3B11 (1:500 Synaptic Systems Mouse – 147111), VGAT (1:1000 Synaptic Systems Rabbit – 131003), and RIM1 (1:500 Synaptic Systems Rabbit – 140003) staining was performed in blocking solution for 1 hour followed by 5 min washes (3X). Coverslips were incubated with appropriate secondary antibodies in blocking solution for 1 hr at RT (for 3D-SIM; 1:1000 ThermoFisher Alexa Fluor 488, 568 and 647) or overnight (for dSTORM; 1:1000 Alexa Fluor 488 anti-rabbit, 1:1000 Sigma-Aldrich CF568 anti-mouse, 1:1000 Abcam Alexa Fluor 647 anti-guinea pig) and (for STED; 1:200 Abberior STAR 580 anti-guinea pig, 1:200 Abberior STAR 635 anti-rabbit, 1:200 Abberior STAR 460L anti-mouse). Coverslips were washed for 5 min (3X) before being mounted on microscope slides with ProLong Gold mounting media (Thermofisher) for 3D-SIM and STED. For dSTORM, coverslips were washed and post fixed with 4% PFA in 1X PBS for 5 min at RT followed by final washes where they remained in 1X PBS until imaging.

#### Immunohistochemistry

As previously described (2, 3), 9-week-old male mice were anesthetized and transcardially perfused with 0.9% saline followed by 4% paraformaldehyde (PFA). Whole brains were extracted and rinsed in 0.1 M phosphate buffer (PB), followed by transfer to a glycerol and Sorenson's Buffer cryoprotection solution to dehydrate the tissue. Frozen coronal sections (50 $\mu$m) were cut using a freezing sliding microtome and placed in a cryostorage solution containing phosphate buffer, ethylene glycol, polyvinylpyrolidine, and sucrose. Free floating sections were washed with 0.1 M PB for 10 min (2X) then with Tris buffer saline (TBS) for 10 min (3X), followed by permeabilization and block (10% NGS and 0.2% TritonX in TBS) for 1 h at RT. Sections were incubated with $GABA_AR$-$\beta$3 (1:500 NeuroMab Mouse 75149) and VGAT (1:500 Synaptic Systems Rabbit – 131003) antibodies in an antibody dilution solution (2% NGS and 0.2% TritonX in TBS) overnight at 4°C on a rocker. Sections were washed for 10 min (3X) in TBS at RT and then incubated with appropriate secondary antibodies (1:200

Abberior STAR 460L anti-mouse, Abberior STAR RED anti-rabbit) for 2 hrs in antibody dilution solution at RT. Prior to mounting with ProLong Glass, sections were washed for 10 min (3X) in TBS followed by 10 min in 0.1 M PB.

### Image Acquisition and Analysis

#### 3D-SIM Imaging

A Nikon SIM-E Structured Illumination super-resolution microscope equipped with a 100x, 1.49 NA objective was used to acquire multichannel SIM images as previously described (1, 2, 4, 5). Illumination from 488, 561, and 640 laser diodes excite the sample and emission light was detected by an ORCA-Flas 4.0 sCMOS camera (Hamamatsu). Nikon Elements software was used for image acquisition. To optimize for a high signal to noise ratio, reduce photobleaching, and to keep the overall count-level below 12000, acquisition conditions and camera integration time were adjusted similarly as in (1). Hippocampal neurons with pyramidal morphology were imaged. The soma, proximal and distal dendrites of each neuron were imaged in sequential acquisitions.

#### 3D-SIM Analysis

Typically, 10-15 ROIs (individual synapses) were manually selected per region of the neuron (4-5 neurons), with a biological replicate of at least 3 neuronal cultures. Selected synapses were positive for VGAT, gephyrin and $GABA_AR$-$\gamma$2, and completely encompassed within the Z-stack. To quantitively characterize synaptic architecture and nanoscale organization, we utilized a high throughput analysis pipeline which processes selected synapses by background subtraction (ImageJ), image segmentation (split-Bregman/MOSAIC suite), and geometric analysis (MATLAB) as previously described (1). For image segmentation, the following parameters were utilized: 'Subpixel segmentation', 'Exclude Z edge', Local intensity estimation 'Medium', Noise Model 'Gauss'. SIM data was transformed into 3D renderings using the function CaptureFigVid.m.1.

#### dSTORM Imaging

Samples for dSTORM were prepared as previously described (2, 4, 5), and imaging was performed on a Zeiss Elyra P.1 TIRF microscope using a Zeiss alpha Plan Apochromat TIRF 100x/1.6 NA oil objective, a tube lens providing an additional factor of 1.6 magnification, and a quad-band dichroic (405/488/561/642). Alexa647 and CF568 dyes were captured by an Andor iXon+ EMCCD camera in sequential time-series of 20,000 frames each at a gain setting of 100 with an integration time of 18ms. Image size was 256 x 256 pixels, with a pixel size of 100 nm xy. Alexa647 molecules were ground-state depleted and imaged with a 100 mW 642 laser at 100% AOTF transmission in ultra high-power mode, corresponding to approximately 1.4W/cm$^2$. Emission light passed through a LP 655 filter. CF568 molecules were ground-state depleted and imaged with a 200 mW 561 laser at 100% AOTF transmission in ultra high-power mode, corresponding to approximately 2.5 W/cm$^2$. Emission light was passed through a BP 570-650 + LP 750 filter. Ground-state return was prompted by continuous illumination with a 50 mW 405 laser at 0.01 to 0.1% AOTF transmission for each dye.

#### dSTORM Processing

dSTORM processing is fully described in (4). Briefly, localization of dye emitters was performed using the ThunderSTORM ImageJ plugin (6). Image filtering was done with the Wavelet filter setting, with a B-Spline order of 3 and scale of 2.0. A first pass approximate localization of molecules was achieved by finding local maximum with a peak intensity threshold of 3*std (Wave.F1) and 8-neighborhood connectivity. Sub-pixel localizations were achieved by weighted least-squares fitting of the PSF by use of an integrated Gaussian with a fitting radius of 4 pixels and an initial sigma of 1.2. Filtering of localizations were based on the attributes of uncertainty (< 20 nm), sigma (50 – 150 nm), and intensity (< 15,000 for Alexa647 and < 10,000 for CF568). Localizations within 50 nm were merged with a frame-gap allowance of 1. Channel registration was performed using custom-written MATLAB scripts and was based on multi-colored bead acquisitions taken at the start of each experiment. Drift correction was performed using the built-in cross-correlation based method in the ThunderSTORM plugin. Localizations were rendered into images using the ThunderSTORM visualization module using the method of average shifted histograms with a magnification of 10 and lateral shift of 2 nm.

**dSTORM Analysis**

Individual synapses were manually selected from a composite SR/WF image and ROI coordinates were recorded with a custom ImageJ macro. The gephyrin scaffold and $GABA_AR$ localizations were segmented using a coordinate-by-coordinate density calculation. Since labeling density can vary greatly, the thresholding parameter was determined from the overall density range of the ROI. Localizations with a local density in the lower 10% of that range were considered outside of the synaptic region. The boundaries for these regions were delineated using the alphaShape function in MATLAB, with an $\alpha$ value of 100. Only regions with an area of 1.5e3 $nm^2$ or greater were considered for analysis.

Based on the methodology used in (7), SSDs were defined by a cutoff determined by randomizing the experimental localizations assuming a uniform distribution across the synaptic region. The local density threshold for an experimental coordinate to be considered as part of an SSD was set at the mean local density of the randomized dataset plus 2 standard deviations. The geometric boundaries of individual SSDs were delineated using the alphaShape function, with an $\alpha$ value of 7. $GABA_AR$-$\gamma$2 SSDs were classified as overlapping with gephyrin SSDs if the overlap area had a fraction of 0.23 or greater of the $GABA_AR$-$\gamma$2 area (similar to (5)).

**3D-STED Imaging**

For imaging in culture, STED super-resolution images were acquired using an Abberior STEDYCON addition to an Olympus confocal microscope. The microscope was equipped with the following: an Olympus x100 STED objective (UPLXAPO100XO, 1.45 NA), 4 excitation lasers; 405 (cw), 485 nm (pulsed), 561 nm (pulsed), 640 nm (pulsed), 4 corresponding single-photon counting APD detectors (avalanche photodiodes), and a 775 nm laser for stimulated emission depletion. Each image was acquired using a minimum of 3 z-stacks at 0.15 $\mu$m spacing and deconvolved using the SVI Huygens deconvolution software with the Standard Mode Express Deconvolution.

For imaging in hippocampal CA1 brain slices, an Abberior Infinity 3D STED microscope was used. The STED system is incorporated onto an Olympus BX63F upright confocal microscope with a V380FM Luigs and Neumann motorized stage. It is capable of wide-field LED illumination and has a quad band filter cube (DAPI/STAR 580/STAR RED) to support imaging with pulsed excitation lasers at 485 nm, 561 nm and 640 nm. For STED, the system has a STED laser module (775-nm pulsed high-power variant). Detection is mediated by three-channel tunable spectral APD detectors and a MATRIX detector to enable background subtraction during STED acquisition. Each optical section was subjected to Huygens deconvolution before image analysis.

**3D-STED Analysis**

Image segmentation was performed using the same pipeline as described in the 3D-SIM Imaging and Analysis methods section, with the following changes to the image segmentation Bregman/MOSAIC suite (8) parameters to better match the data: Local intensity estimation 'High', Noise Model 'Poisson'. The alignment between RIM1 and $GABA_AR$ was determined by methods previously described using randomized data (5). For the RIM1/$GABA_AR$ pairing analysis we identified the maximum intensity point within each SSD. The binary mask created by the object-segmentation was applied to the original intensity image, creating an object-bound intensity map. The center points of regional intensity maximum were then identified using the MATLAB function [imregionalmax]. The nearest neighbor regional intensity max point pairs in the corresponding channels (RIM1 and $GABA_ARs$) were assigned based on the [knnsearch] function in MATLAB. To ensure realistic synaptic geometries and object numbers, randomized data for coupling analysis was generated on a ROI-to-ROI basis. The synaptic compartments were used to create a bounding box for the randomized points, the number of which were set to match the number of objects in the experimental data for that ROI (1, 5).

**Computational Modeling of Inhibitory Synaptic Currents**
Monte-Carlo simulations were performed using the mCell platform (version 4.0.6) (9, 10). Model geometries based on our STED/SIM data, were constructed in the CellBlender (version 2.93) module of mCell. Each synapse type consists of a pre- and post-synaptic membrane (2.5 $\mu m^2$) with reflective surface properties and a cleft distance of 30 nm. The diffusion coefficient of GABA was set at 49 $\mu m^2$sec-1 (11). Simulations consisted of 100,000 time-points of 10-6 seconds per time-point. GABA release occurred synchronously at time-point zero. 500 GABA molecules were released from each release site (11, 12). Each model simulation was run 160 times. Receptor reaction parameters were based on the model by Jones and Westbrook (13), with minor modifications: since the open fraction of channels plateaued at ~ 0.20 (see (12) for similar traces), we allowed for GABA to disassociate from both open and desensitized receptors; the original schema only allowed for GABA disassociation from closed channels, creating a kinetic bottleneck that trapped most the receptors in a dually bound state. The same GABA kon, koff rates were used for all three channel states.

IPSCs were calculated using the formula:

$$I(t) = [g \times n(t)] \times (Vm - Vr)$$

Where *g* is the signal channel conductance, 27 psec. *n(t)* is the number of open GABA receptors at time-point *t*, *Vm* is the resting membrane potential (-75 mV) while *Vr* is the reversal potential (-85 mV) (14).

GABA release-sites and receptor SSD pairs were classified as follows: *Aligned*, lateral center-center distance <= 20 nm; *Offset*, lateral center-center distance ~20-50 nm; *Orphan*, nearest SSD center > 100 nm; *Uniform*, uniform distribution of the receptor across the synapse with no SSD clustering.

The following synapse-specific parameters we used for Fig. 5:

| *Dendritic synapse* | **Fig. 5 C-F** |
|---|---|
| PSD area | 400 $nm^2$ |
| Total receptor # in PSD | 80 |
| Receptor # per SSD | 19 |
| $GABA_AR$ SSD area | 47 $nm^2$ |

| | |
|---|---|
| $GABA_AR$ SSD # | 2 |
| Release site # | 2 |
| Trans-synaptic arrangement | Aligned (5C, F); Offset (5D); Uniform (5E). |

| *Dendritic synapse (Large)* | **Fig. 5 G, H** |
|---|---|
| PSD area | 500 $nm^2$ |
| Total receptor # in PSD | 134 |
| Receptor # per SSD | 27 |
| $GABA_AR$ SSD area | 55 $nm^2$ |
| $GABA_AR$ SSD # | 3 |
| Release site # | 3 |
| Trans-synaptic arrangement | Aligned (5G); Uniform (5H). |

| *Somatic synapse (small)* | **Fig. 5 I** |
|---|---|
| PSD area | 400 $nm^2$ |
| Total receptor # in PSD | 96 |
| Receptor # per SSD | 27 |
| $GABA_AR$ SSD area | 55 $nm^2$ |
| $GABA_AR$ SSD # | 2 |
| Release site # | 2 |
| Trans-synaptic arrangement | Aligned (5I) |

| *Somatic synapse (large)* | **Fig. 5 J-L** |
|---|---|
| PSD area | 500 $nm^2$ |
| Total receptor # in PSD | 134 |
| Receptor # per SSD | 27 |
| $GABA_AR$ SSD area | 55 $nm^2$ |
| $GABA_AR$ SSD # | 3 |
| Release site # | 3 |
| Trans-synaptic arrangement | 3 aligned + 1 orphan (5J); Uniform (5K); 1 aligned + 3 orphan (5L). |

**Statistical Analysis**

Statistical analysis was performed using Graphpad Prism 11 software. Data was assessed for normality using both the D'Agostino-Pearson test and the Shapiro-Wilk test. Non-parametric data was evaluated using the Mann-Whitney test for two experimental conditions or the Kruskal-Wallis with Dunn's Test for more than two conditions. Parametric data was analyzed using the Unpaired Student's t-test (two conditions) or the One-Way ANOVA with Dunnett's test (> 2 conditions). Statistical analysis of paired data used the paired t-test. To test for linear correlation, the 'simple linear regression' analysis in Prism was used without constraints to determine if the slope deviated significantly from zero and if the slopes of multiple conditions were different. For all statistical analyses p values $< 0.05$ were considered significant.
